# Identification of ERAD-dependent degrons for the endoplasmic reticulum lumen

**DOI:** 10.1101/2023.06.21.546000

**Authors:** Rachel Sharninghausen, Jiwon Hwang, Devon D. Dennison, Ryan D. Baldridge

## Abstract

Degrons are minimal protein features that are sufficient to target proteins for degradation. In most cases, degrons allow recognition by components of the cytosolic ubiquitin proteasome system. Currently, all of the identified degrons only function within the cytosol. Using *Saccharomyces cerevisiae*, we identified the first short linear sequences that function as degrons from the endoplasmic reticulum (ER) lumen. We show that when these degrons are transferred to proteins, they facilitate proteasomal degradation through the ERAD system. These degrons enable degradation of both luminal and integral membrane ER proteins, expanding the types of proteins that can be targeted for degradation in budding yeast and mammalian tissue culture. This discovery provides a framework to target proteins for degradation from the previously unreachable ER lumen and builds toward therapeutic approaches that exploit the highly-conserved ERAD system.

## Introduction

Protein degradation plays an essential role in regulating diverse cellular processes including cellular signaling, metabolic adaptation, and cell cycle regulation. The ubiquitin proteasome system is the primary cellular degradation route, accounting for over 80% of protein degradation^1^. Ubiquitination requires the concerted action of the ubiquitination cascade comprising E1 ubiquitin activating enzymes, E2 ubiquitin conjugating enzymes, and E3 ubiquitin ligases. For ubiquitin, in *S. cerevisiae* there is one E1, eleven E2s, and >60 E3s whereas mammals have two E1s, ∼40 E2s, and >600 E3s^2^. The specificity of the ubiquitination process is primarily driven by the E3 ubiquitin ligases and a major challenge in ubiquitin biology is the identification of the sequences that target proteins for degradation (called “degrons”).

Degrons are usually short linear motifs and, by definition, are sufficient to confer degradation when transferred to otherwise stable proteins. Degrons can be acquired, or inherent. Acquired degrons are generally post-translational modifications that can be based on proteolytic cleavage, phosphorylation, or acetylation. Inherent degrons are features of the primary polypeptide sequence formed by linear or conformational epitopes. Inherent degrons can be shielded when proteins are appropriately-folded or incorporated into larger protein complexes. The first degrons to be discovered were at the amino-terminus of proteins^3^ and these N-degrons were eventually summarized as the “N-end rule”^4^, and later the “N-degron pathways”^5^. Recent systems-level analyses have expanded the availability of known degrons broadly^6,7^, with a series of “C-end rules”^8,9^, and additional variations of the N-degron pathways (reviewed in ^10^).

Even with a wide range of physiological roles for protein degradation, degrons are still unidentified for most E3 ubiquitin ligases. All known degrons target cytosolic proteins for ubiquitination and degradation by the proteasome and the identification of degrons for a few key ubiquitin ligases has enabled exploitation of the proteasome to facilitate targeted protein degradation of “undruggable” cytosolic proteins^11^. However, many proteins that originate from the lumen of membrane-encapsulated organelles are also degraded using the proteasome and degrons for these organelles are mysterious^12^. Therefore, targeting proteins for degradation from within organelles requires a detailed understanding of local organellar protein quality control systems.

The endoplasmic reticulum (ER) represents the organelle with the largest flux of proteins, with over 40% of proteins translocated into the ER before trafficking to other organelles or secretion from the cell. Both soluble luminal proteins and integral membrane proteins are folded in the ER and undergo quality control before being released into the secretory pathway. At the ER, the primary protein quality control pathways are, collectively, referred to as endoplasmic reticulum associated degradation (ERAD) and these systems are highly conserved among all eukaryotes^13–18^. One such system, the Hrd1-centric ERAD complex, recognizes proteins not passing quality control and retrotranslocates them from the ER lumen to the cytosol for ubiquitin-mediated proteasomal degradation. In *Saccharomyces cerevisiae,* the Hrd1 complex comprises 5 proteins: Hrd1, Hrd3, Usa1, Der1, and Yos9. In the cytosol, this pathway requires a highly conserved AAA-ATPase (Cdc48), its cofactors (Ufd1 and Npl4), and the ubiquitination proteasome system to degrade ERAD substrates^19–22^. Soluble, luminal ERAD substrates are retrotranslocated by hetero-oligomers of Hrd1/Der1^23–25^, or in some cases, homo-oligomers of Hrd1^26–28^. Hrd1 is sufficient for the basic functions of ERAD but without the other complex components, loses the specificity that normally defines the system^14^. This system has broad specificity and seems to be able to distinguish folded and unfolded proteins^29^. Despite nearly 3 decades of study, degrons (neither sequences nor features) that allow degradation through the Hrd1-centric ERAD pathway remain a complete mystery^30^.

Here, we have identified the first short luminal degrons of the Hrd1-centric ERAD pathway. We demonstrate that a degron can be functionalized to drive protein degradation from the ER lumen, a previously inaccessible cellular location. We show that this technology can drive degradation of both soluble, luminal ER proteins and integral membrane ER proteins. This degron works in both budding yeast and mammalian tissue culture. This work provides an exciting and simple method of targeting proteins for degradation from within the ER by exploiting the highly conserved ERAD system.

## Results

### Identification of ER-localized degrons

The ER is the primary location for protein quality control within the secretory pathway, but how ER protein quality control systems distinguish folded from unfolded proteins is unclear. We wanted to understand the degrons recognized within the ER lumen and started by designing an ER-targeted reporter of protein stability. We targeted a tandem fluorescent protein timer (tFT) to the ER to function as a reporter of protein stability (ER-tFT, Figure 1A and S1A). tFTs contain a fast-maturing fluorescent protein (here, superfastGFP^31^) and a slower-maturing fluorescent protein (here, mCherry^32^) and have been used effectively to identify N-terminal degrons that function in the cytosol and nucleus^33,34^. By measuring the ratio of mCherry to GFP fluorescence, a protein’s stability can be assessed; the lower the ratio, the more unstable the protein. To test whether the ER-tFT could successfully distinguish stable from unstable proteins, we compared the ER-tFT to a well-characterized, unstable, luminal ERAD substrate, KHN, tagged with the tFT (KHN-tFT, Figure 1A). Using a cycloheximide chase followed by immunoblotting, we found the ER-tFT was quite stable, while the KHN-tFT was degraded with a half-life of less than 30 minutes (Figure 1B), consistent with previous reports^35^. Using flow cytometry, the two proteins were also distinguishable following cycloheximide treatment (Figures 1C, 1D and S1B). After establishing the tFT reporter could distinguish protein stability within the ER, we turned our attention to identifying luminal degrons.

**Figure 1.**
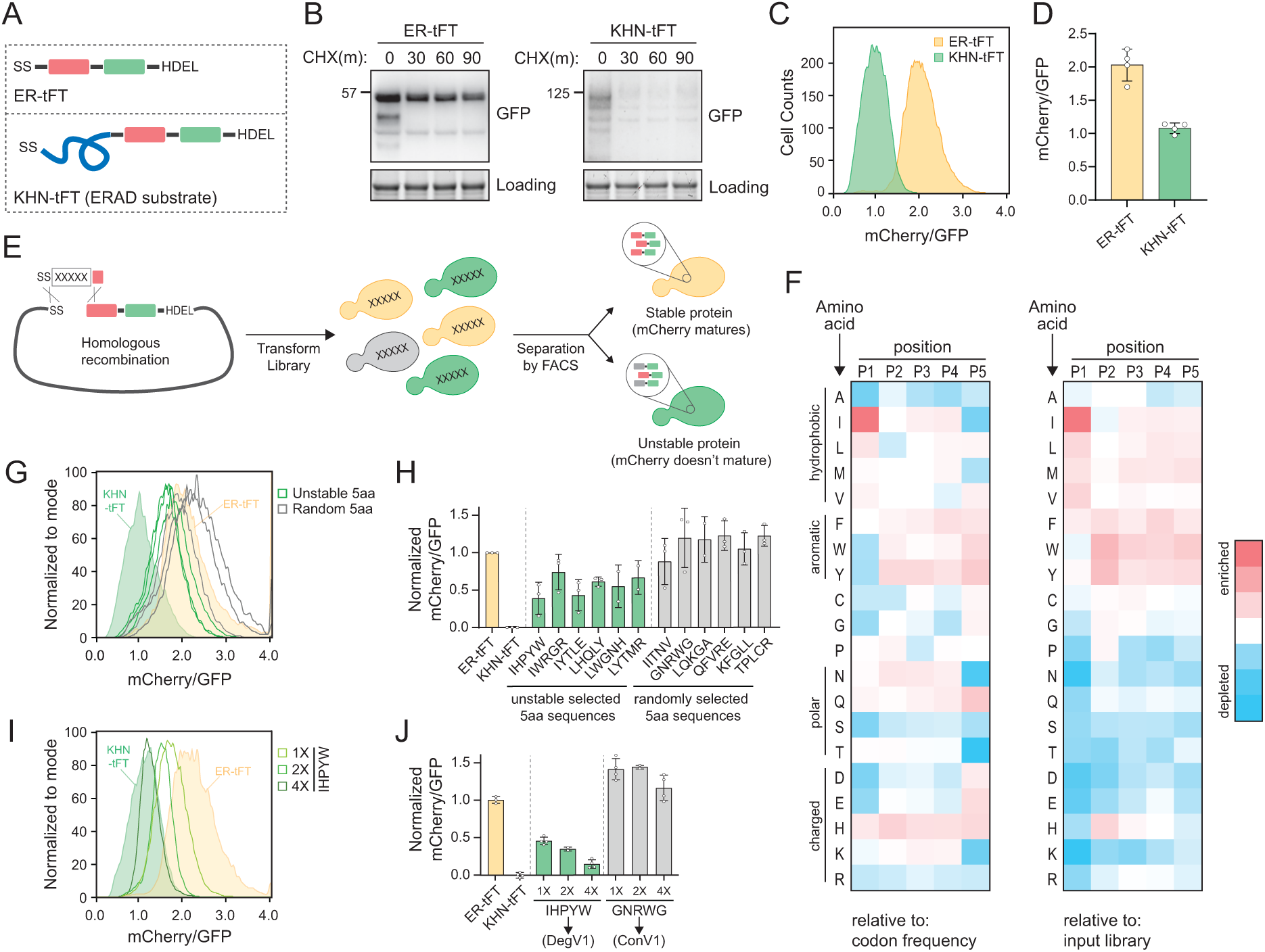
Identification of endoplasmic reticulum localized degrons. A) Schematic depicting the ER-tFT and KHN-tFT constructs, which contain an ER-targeting signal sequence (SS), mCherry (red), superfastGFP (green), and the HDEL ER-retention sequence. KHN-tFT functions as a quickly degraded ERAD substrate (a positive control for degradation). B) Wild-type yeast expressing the constructs described in (A) were treated with cycloheximide (CHX) for 0, 30, 60, or 90 minutes, harvested, and protein levels were assessed by immunoblotting using anti-GFP antibodies. Total protein was visualized in gel using stain-free technology (Loading). C) Flow cytometry of yeast expressing the constructs in (A) treated with CHX for 2 hours. The mCherry/GFP fluorescence intensity ratio of each cell was calculated and plotted. D) Quantification of the mean mCherry/GFP ratio of four biological replicates as in (C). E) Overview of the pentapeptide library generation and isolation of unstable variants using fluorescence activated cell sorting (FACS). A DNA fragment containing the pentapeptide-ER-tFT library was electroporated with linearized ER-tFT plasmid. The resulting yeast library contains a mixture of variants that are separated by FACS, with less stable variants having decreased mCherry/GFP fluorescence intensity compared to stable variants. F) Heatmap of amino acid enrichments at each position within the unstable pentapeptide library. Values are displayed relative to either codon usage (left) or relative to the input library (right). G) As in (C) with strains expressing either ER-tFT (yellow fill), KHN-tFT (green fill), individual FACS isolates from the “unstable” pentapeptide sequences (green lines), or randomly selected pentapeptide-ER-tFT sequences from the input library (gray lines). H) Quantification of at least 2 biological replicates conducted as in (G). The unstable groups (green) and random groups (gray) were significantly different from each other using a one-way ANOVA and Tukey’s multiple comparisons tests. I) As in (C) with strains expressing either ER-tFT (yellow fill), KHN-tFT (green fill), a single IHPYW (1X), 2x repeat of IHPYW (2X), or 4x repeat of IHPYW at the N-terminus of ER-tFT. J) Quantification of 2-3 biological replicates of (I).

To identify degrons that function within the ER lumen, we generated a library of short, linear peptide sequences embedded into the ER-tFT. Using PCR with degenerate primers, we generated an unbiased pentapeptide library encoded in a DNA fragment to use for homologous recombination in cells (Figures 1E, S1C, and S1D). The theoretical amino acid diversity of a pentapeptide library is 3.2 million (20^5). We transformed the library into wild-type yeast and obtained 1.2 million transformants. Using fluorescence-activated cell sorting (FACS) we separated cells expressing the pentapeptide-ER-tFT by their mCherry/GFP ratio (Figure S1E and S1F). We collected an “unstable” bin (exhibiting a low mCherry/GFP ratio), which encompassed 4% of the sorted cells, for sequencing (Figure S1F).

The sequencing results of the unstable sorted bin illuminated an enrichment of pentapeptides beginning with isoleucine and leucine (Supplementary data S1). Specifically, isoleucine at the first position was present in 17.7% of unstable pentapeptides while leucine was present in 14.8%. Based on codon usage in a random sampling, isoleucine was predicted to appear 4.7% of the time (3/64 codons) giving a 3.8 fold enrichment in our dataset. Leucine was predicted to appear at a specific position 9.4% of the time (6/64 codons) giving a 1.6 fold enrichment in our dataset (Figure 1F). When compared to the input library abundance, enrichment corresponds to 2.7 and 1.5 fold, respectively (Figure 1F). Rather than attempting to build a consensus sequence that would have to report on many steps in ER quality control, we selected pentapeptide sequences present in the unstable bins that broadly represented the enrichment trends we observed to individually clone and characterize (Figure 1F and Supplementary data S1).

We compared the stability of six different pentapeptides with isoleucine or leucine at position one to ER-tFT alone, to KHN-tFT, and to a set of randomly selected pentapeptides. KHN-tFT was the least stable, followed by pentapeptides selected from the unstable bin and containing isoleucine or leucine at position one. Several of the randomly selected pentapeptides exhibited a slight reduction in stability, relative to ER-tFT alone, but pentapeptides selected from the unstable bin were consistently less stable (Figure 1G, 1H, and S1G). Encouraged by our results, we sought to find a peptide sequence that would match KHN-tFT instability. We found that repeats of IHPYW, one of the most unstable sequences, dramatically decreased protein stability, with a 20 amino acid 4x(IHPYW) repeat successfully resembling the KHN control (Figure 1I and 1J). On the other hand, simply repeating a stable control sequence (GNRWG) did not destabilize the protein (control variant 1 (ConV1), Figure 1J). Based on these results, we concluded that the 4x(IHPYW) sequence, which we called DegV1 (<u>Deg</u>ron <u>V</u>ariant 1), functions as an ER-localized degron.

### DegV1 is an ERAD-dependent degron degraded by the proteasome

We next tested whether DegV1 functioned as an ER luminal degron by transferring it from the original ER-tFT reporter (Figure 1) to a second protein known to fold in the ER lumen. We targeted the LaG16 anti-GFP nanobody^36^ to the ER using the mating factor alpha signal sequence and an ER retention signal (ER-NbGFP, Figure S2A). First, to ensure the signal sequence was removed and did not contribute to the DegV1 degron, we compared the molecular weight of a cytosolic NbGFP (without signal sequence), a cytosolic DegV1-NbGFP (without signal sequence), and the ER-DegV1-NbGFP (with signal sequence)(Figure 2A). Based on the size of the ER-DegV1-NbGFP compared to DegV1-NbGFP, we concluded that ∼90% of the ER-targeted construct had the signal sequence removed and uncleaved signal sequence was unlikely to factor into the function of DegV1 (Figure 2A). Using a cycloheximide chase, the ER-NbGFP protein alone was quite stable, but embedding DegV1 after the N-terminal signal sequence destabilized ER-NbGFP with a half-life of approximately 30 minutes (Figure 2B). This result validated DegV1 as the first, relatively short, degron sequence that can be used for targeting ER luminal substrates for degradation. Next, we tested if DegV1 could also function as an internal or C-terminal degron. To test whether the DegV1 sequence worked within internal loops, we used the same nanobody scaffold but replaced the complementary determining region 3 with DegV1 and found that the presence of an internal DegV1 made no difference in protein stability when compared to the nanobody alone (Figure 2C and S2A). Testing DegV1 at the C-terminus of the NbGFP was complicated by the requirement for an ER retention signal. We integrated the DegV1 sequence immediately before the HDEL retention signal, and, again, this resulted in no change in the stability of the NbGFP target protein (Figure 2D and S2A). The ER retention signal is an important component of this construct’s innate stability, because loss of the HDEL signal, with or without the DegV1 sequence at the extreme C-terminus, results in degradation in the vacuole or secretion from the cell (Figure 2D). Together, these results suggest that the DegV1 degron was capable of acting as an ER luminal degron, but only when positioned at the N-terminus of ER-localized proteins.

**Figure 2.**
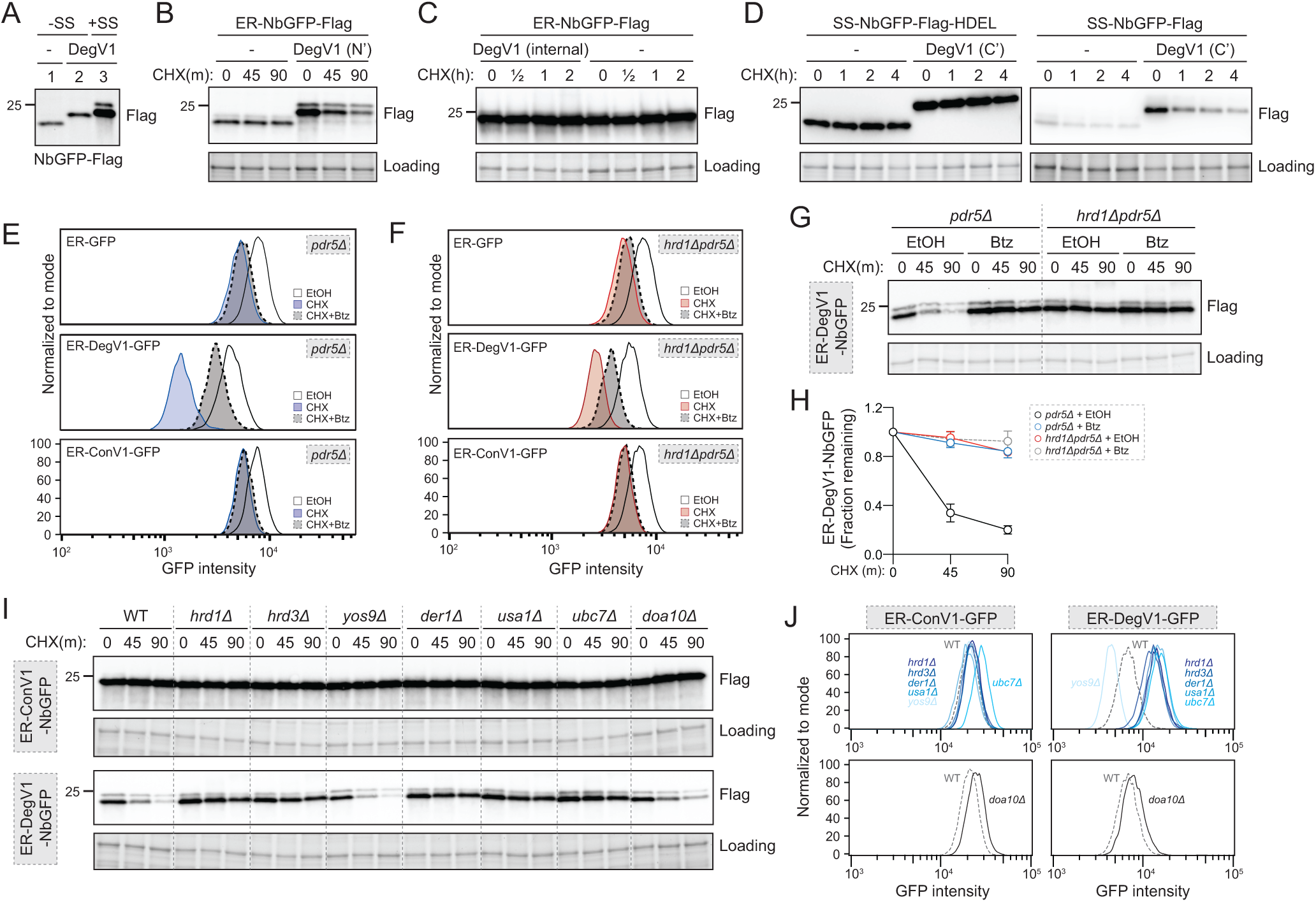
DegV1 is an ERAD-dependent degron degraded by the proteasome. (A) The apparent molecular weight of anti-GFP nanobodies were assessed (NbGFP-Flag) either with or without signal sequence, in the presence or absence of DegV1, were assessed by SDS-PAGE electrophoresis followed by immunoblotting with anti-Flag antibody. This panel is representative of two biological replicates. (B) The degradation of ER-targeted anti-GFP nanobodies (ER-NbGFP-Flag) either with, or without, DegV1 were monitored following addition of cycloheximide (CHX), using SDS-PAGE and immunoblotting. Loading controls were visualized by stain-free technology. (C) The degradation of ER-NbGFP-Flag with DegV1 replacing the CDR3 region was analyzed as in (B). (D) The degradation of a nanobody with DegV1 located either directly preceding the C-terminal HDEL ER-retention signal (left) or directly at the C-terminus of the nanobody (right) was analyzed as in (B). (E) Degradation of the ER-targeted proteins ER-GFP (top panel), ER-DegV1-GFP (middle panel), or ER-ConV1-GFP (bottom) were analyzed in a *pdr5Δ* strain by flow cytometry following either ethanol (EtOH) or cycloheximide (CHX) treatment for 2 hours. Where indicated, cells were pretreated with bortezomib (Btz) for 2 hours. (F) As in (E), but in a *hrd1Δpdr5Δ* strain. (G) Degradation of ER-DegV1-NbGFP was followed in *pdr5Δ or hrd1Δpdr5Δ* strain as in (B). Where indicated, cells were pretreated with bortezomib (Btz) for 2 hours. (H) Quantification of (G) with error bars representing the standard deviation. (I) Degradation of ER-ConV1-NbGFP (top panel) or ER-DegV1-NbGFP (bottom panel) was followed in of ERAD component deletion strains as in (B). This panel is representative of two independent biological replicates. (J) The degradation of ER-ConV-GFP (left), or ER-DegV-GFP (right) were analyzed in the indicated ERAD component deletion strains by flow cytometry following cycloheximide treatment for 2 hours. All panels in this figure are representative of at least three independent biological replicates, unless otherwise indicated.

DegV1 was able to actively target proteins for degradation from the ER lumen. The majority of protein degradation occurs at the proteasome, so we next tested whether DegV1-tagged proteins were degraded by the proteasome. We confirmed ER-GFP was ER-localized (Figure S2B) and examined the stability of the fluorescent ER-localized construct GFP alone, or with DegV1 or ConV1 in cells treated for 2 hours with the proteasomal inhibitor bortezomib and/or cycloheximide. The stability of ER-GFP alone was similar either in the presence (dashed outline) or absence (solid outline) of bortezomib (panel 1, Figure 2E). Appending DegV1 immediately after the signal sequence resulted in degradation of ER-GFP (solid blue line), which was inhibited by adding bortezomib (dashed line, panel 2, Figure 2E). This indicated DegV1 targets the luminal ER-GFP for proteasomal degradation. As expected, appending a control sequence (ConV1) of the same length was similarly stable to ER-GFP alone and stability was not affected by bortezomib (solid line versus dashed line, panel 3, Figure 2E, see also S2C). Therefore, DegV1-targeted luminal ER substrates were degraded by the proteasome.

Proteasomal degradation of luminal ER proteins is mediated by the Hrd1-ERAD system^37^. Consequently, we suspected that Hrd1-centered ERAD mediates DegV1-targeted proteasomal degradation. Again, we tested the stability of ER-GFP in cells lacking the central component to the Hrd1-ERAD system, the ubiquitin ligase Hrd1. The steady state levels of ER-GFP alone or with ConV1 were similar in the absence of Hrd1 and remained unaffected by proteasome inhibition (panels 1 and 3, Figure 2F). In contrast, DegV1-containing ER-GFP was more stable in a *hrd1Δ* strain, and bortezomib resulted in little further stabilization of DegV1-containing GFP (panel 2, Figure 2F, see also S2D), possibly from a small fraction being incompletely translocated into the ER. In the absence of Hrd1, we observed some ER leakage of ER-DegV1-GFP to the vacuole and appearance of a degradation resistant GFP fragment that resembled the known luminal ERAD substrate, CPY*-GFP (indicated by * in Figure S2E and S2F). Altogether, these results are consistent with a role for Hrd1 in the degradation of ER-DegV1-GFP.

To further test the role of Hrd1 in DegV1-targeted degradation of luminal ER substrates, we tested additional soluble ER proteins (Figure 2G and S2G) that were, otherwise, relatively stable in the ER lumen (Figure S2G). As expected, when DegV1 was appended to the ER-NbGFP, we found the protein was unstable, with a half-life of approximately 30 minutes in a cycloheximide chase (Figure 2G, 2H, and S2G). Next, we tested whether other known components of the Hrd1 ERAD complex were required for DegV1 degradation. We found that, with the notable exception of Yos9, the previously-identified components of the ERAD-L complex (Hrd1, Hrd3, Der1, and Usa1) were required for DegV1 degradation (Figure 2I and 2J). Conversely, the Doa10 ubiquitin ligase was not required (Figure 2I and 2J). Consistently, degradation of each of DegV1-containing proteins were inhibited by either treating cells with bortezomib or genetic deletion of Hrd1.

When DegV1-containing proteins were targeted to the ER lumen, Hrd1-ERAD was required for their degradation (Figure 2E-2J). We tested whether DegV1 would also function as a degron in the cytosol by removing the signal sequence. Somewhat surprisingly, we found that DegV1 also mediated proteasome-dependent degradation when localized to the cytosol (Figure S3). However, in contrast to ER localized DegV1, when DegV1-containing proteins were localized to the cytosol, Hrd1 was not required for their degradation through the proteasome (Figure S3).

In these experiments, we confirmed that DegV1 targets heterologously expressed ER luminal proteins for ERAD-mediated proteasomal degradation. We next tested whether DegV1 could target endogenous *S. cerevisiae* proteins for degradation. We transplanted DegV1 onto three different classes of endogenous, ER-localized proteins. First, we used the endogenous protein Suc2. We used the Suc2 signal sequence followed by either DegV1, or ConV1, an HA tag, the Suc2 coding sequence, a Flag tag, and an HDEL (Figure 3A). With ConV1, Suc2 was stable over several hours, and, based on the modified glycosylation pattern, even appeared to be partially trafficked from the ER. In contrast, with DegV1 Suc2 was dramatically destabilized (Figure 3A). Next, we attached DegV1 to a type I integral membrane ER protein, called Big1, that has its N-terminus in the ER lumen and contains a single transmembrane segment^38^. We replaced the signal sequence of Big1 with the signal sequence of mating factor alpha followed by either DegV1 or ConV1, an HA tag, and the Big1 coding sequence. DegV1 was also able to destabilize the integral membrane protein Big1 (Figure 3B). Finally, we attached DegV1 or the control sequence to the N-terminus of Elo1, a multi-spanning integral membrane protein with 7 probable transmembrane segments and the N-terminus in the ER lumen^39,40^. We found that DegV1 was capable of driving degradation for the multi-spanning membrane protein Elo1 (Figure 3C). These results support that DegV1 functions as an N-terminal degron for endogenous proteins with a range of topologies.

**Figure 3.**
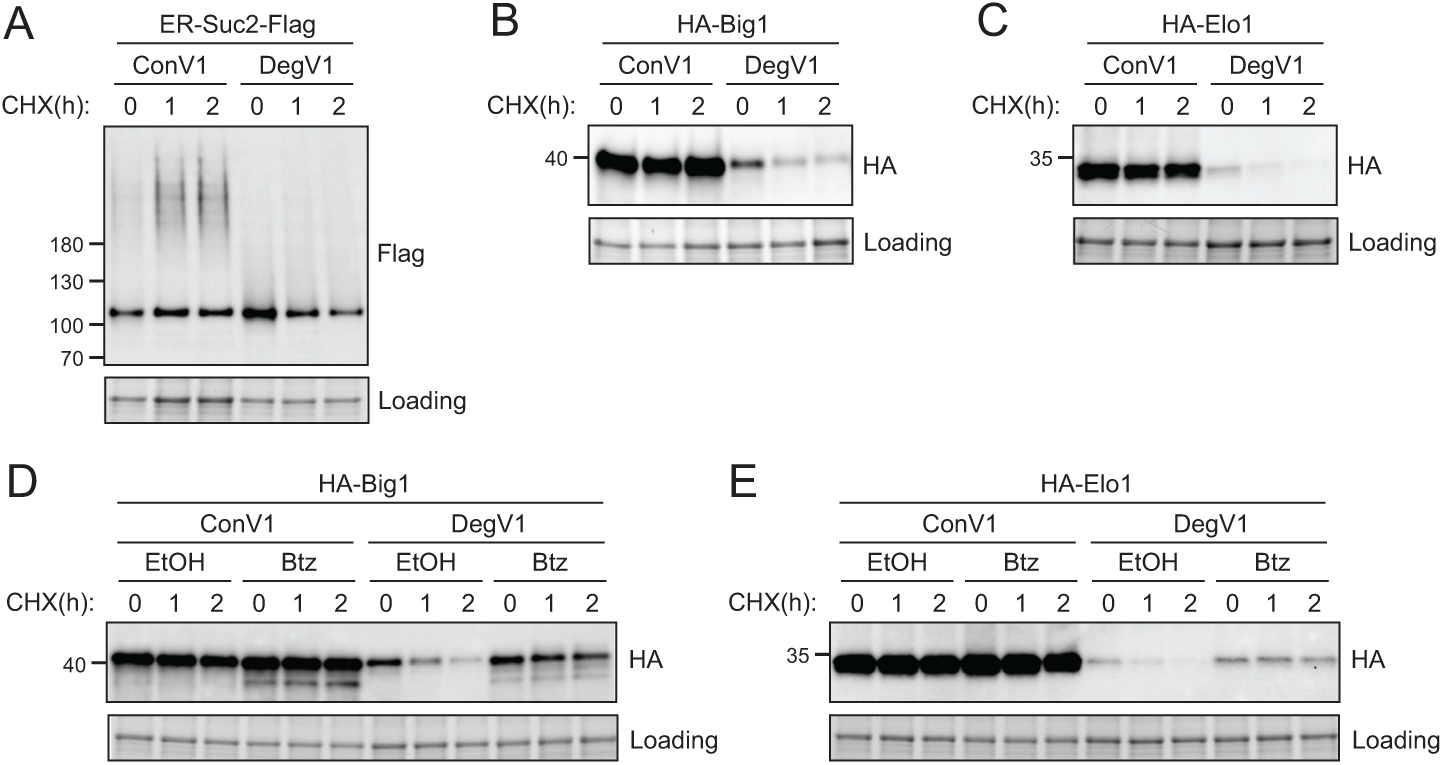
DegV1 targets endogenous ER proteins for degradation. (A) The degradation of an endogenous secretory protein with a C-terminal Flag (ER-Suc2-Flag) containing either DegV1 or ConV1 was monitored following addition of cycloheximide (CHX), using SDS-PAGE and immunoblotting. Loading controls were visualized by stain-free technology. (B) The degradation of a single transmembrane segment ER resident protein (Big1) with the N-terminus in the ER lumen, appended with either DegV1 or ConV1, was followed as in (A). (C) The degradation of polytopic integral membrane ER resident protein (Elo1) with the N-terminus in the ER lumen, appended with either DegV1 or ConV1, was followed as in (A). (D) The degradation of Big1 with DegV1 or ConV1 was followed as in (A) but following a 2 hour pretreatment with either ethanol (EtOH) or bortezomib (Btz) in a *pdr5Δ* strain. (E) The degradation of Elo1 with DegV1 or ConV1 was followed as in (D). All panels in this figure are representative of at least three independent biological replicates.

Soluble, luminal DegV1-containing proteins are targeted to the proteasome by the Hrd1-ERAD pathway (Figures 2E-J). To test whether DegV1-containing integral membrane proteins are also degraded by the proteasome, we followed Big1 and Elo1 degradation after treatment with cycloheximide and bortezomib. Both membrane proteins were significantly stabilized upon treatment with bortezomib (Figure 3D and 3E). Therefore, DegV1 targets both luminal and integral membrane ER proteins for recognition by ERAD and subsequent degradation by the proteasome.

### DegV1 is a functional degron in mammalian cells

DegV1 is a degron facilitating degradation from the ER lumen and also represents the first short, portable degron tag (<180 amino acids^27^) identified for the Hrd1-ERAD system. We turned our attention to the possibility of using DegV1 as a tool in mammalian cells. To determine whether DegV1 functioned as an ER degron in mammalian cells, we generated an ER-targeted mNeonGreen^41^ by appending an N-terminal BiP signal sequence, the HA epitope tag, and the C-terminal ER retention peptide (KDEL) (ER-HA-mNG). We transfected U2OS cells with the ER-mNG containing either ConV1 or DegV1. The addition of DegV1, but not ConV1, reduced the steady-state ER-mNG levels compared to the control (Figure 4A, compare lanes 1, 4, and 7). When we inhibited translation with emetine, we found that ER-mNG and ER-ConV1-mNG were quite stable (Figure 4A and 4B). In contrast, ER-DegV1-mNG was unstable, with a half-life of approximately 4 hours (Figure 4A and 4B).

**Figure 4.**
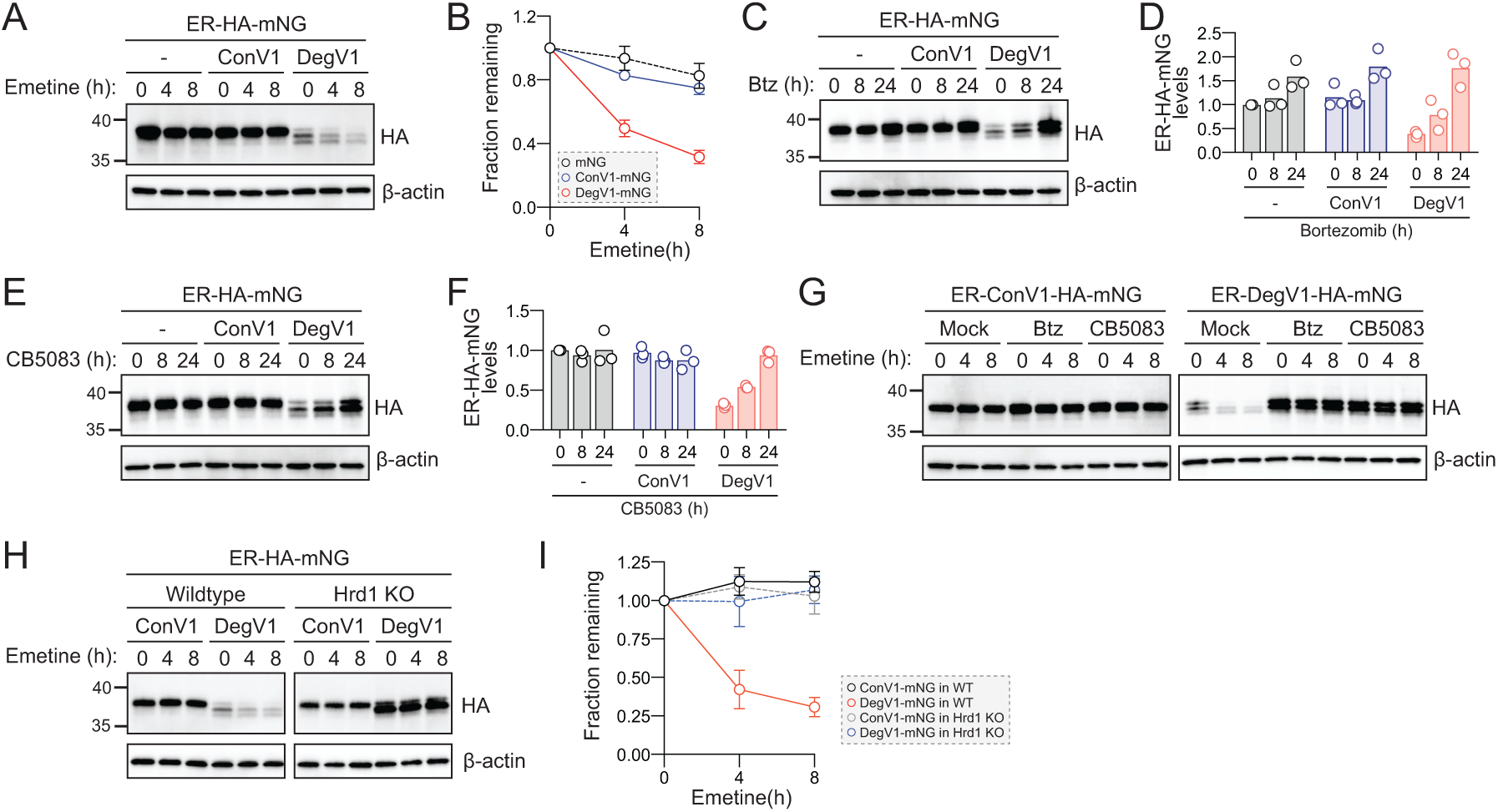
DegV1 functions as a degron in mammalian cells. (A) ER-targeted mNeonGreen (ER-HA-mNG) was expressed alone (-),with ConV1, or DegV1 in U-2 OS cells by transient transfection. The degradation of ER-mNG was followed by immunoblotting with anti-HA antibody after treatment with 50μM emetine. β-actin was used as a loading control. (B) Anti-HA band intensities from (A) were quantified and normalized to the corresponding β-actin level. (C) As in (A) but after treatment with 50nM bortezomib (Btz) for the indicated times. (D) Quantification of (C) normalized to the control protein (ER-HA-mNG). (E) As in (A) but after treatment with 1µM CB5083, a p97 inhibitor, for the indicated times. (F) Quantification of (E) normalized to the control protein (ER-HA-mNG). (G) ER-HA-mNG with either ConV1 (left panel) or DegV1 (right panel) were expressed in U-2 OS pretreated with either 50nM bortezomib or 1µM CB5083 for 16 hours prior to an emetine chase. (H) The degradation of ER-HA-mNG with either ConV1 or DegV1 was followed in HEK293T cells or *HRD1^-/-^* cells using an emetine chase. (I) Quantification of (H). All panels in this figure are representative of at least three independent biological replicates and the quantification is presented as the mean +/-standard deviation.

In *S. cerevisiae*, DegV1 degradation was dependent on the proteasome and mediated through ERAD. In U-2 OS cells, we tested whether degradation of DegV1 was also proteasome-dependent. We treated cells with bortezomib and found that, after an 8 hour treatment, ER-mNG and ER-ConV1-mNG levels remained largely unchanged, but after 24 hours we observed a ∼50% increase. In contrast, for ER-DegV1-mNG proteasomal inhibition resulted in a dramatic accumulation of protein within 8 hours of treatment (Figure 4C and 4D). Remarkably, at 24 hours of bortezomib treatment ER-DegV1-mNG accumulated to similar levels compared to ER-ConV1-mNG, suggesting that the low steady-state level of ER-DegV1-mNG was caused by continuous degradation, rather than a general expression problem.

ERAD-dependent proteasomal degradation also requires the AAA-ATPase p97/VCP (Cdc48 in yeast). Therefore, we tested the stability of the ER-mNG proteins after treatment with the VCP inhibitor CB5083^42^. Similar to the bortezomib treatment, we found that treatment with CB5083 resulted in stabilization of ER-DegV1-mNG (Figures 4E and 4F). As the CB5083 incubation length increased, we found that the levels of ER-DegV1-mNG approached those of the stable, control proteins. To confirm that bortezomib and CB5083 were preventing the active degradation of ER-DegV1-mNG, rather than just improving expression, we analyzed the degradation of the ER-mNG proteins in the presence of bortezomib or CB5083. We confirmed that, without addition of bortezomib or CB5083, ER-ConV1-mNG was stable while ER-DegV1-mNG was degraded (Figure 4G). When we pretreated cells with either bortezomib or CB5083 and inhibited translation with emetine, we found that the degradation of ER-DegV1-mNG was completely inhibited. Finally, we tested whether Hrd1 was required for degradation of ER-DegV1-mNG in mammalian cells. Using either wild-type or Hrd1 knockout cells^43^, we followed degradation of ER-DegV1-mNG and found that the degron-containing protein was stabilized in the absence of Hrd1 (Figure 4H and 4I). Taken together, these data indicate that DegV1 functions as a Hrd1-dependent ER-localized degron in mammalian cells.

## Discussion

In this study, we identified the first short, linear degron motifs that target proteins for degradation from the ER (Figure 1). We focused on functionalizing these motifs and found that DegV1 works with folded, luminal, and completely soluble proteins as well as integral membrane proteins with differing topologies. It works with both exogenous and endogenous proteins that are otherwise stable. This degron appears to only work at the N-terminus of nascent proteins, rather than internally or C-terminally. Importantly, DegV1 is degraded through the Hrd1-ERAD axis using the proteasome (Figures 2 and 3). Furthermore, we found that DegV1 works across eukaryotes in the mammalian cell culture system (Figure 4). Our data highlight a robust, highly potent degron that can be used for targeted protein degradation from the ER lumen and membrane.

Despite a growing understanding of how the ERAD system functions, the fundamental question of how degrons are recognized by ERAD remains unanswered. Here, we identified a number of different degrons, that remain to be completely characterized, but focused on a single degron (DegV1). It is somewhat surprising that DegV1 seems to only work as an N-terminal degron, but it is possible that when DegV1 is positioned anywhere except the N-terminus, DegV1 could be buried within other parts of the proteins. Although this position-dependent effect is simply a result of the methods used to identify this particular degron, it highlights the importance of future work to identify degrons capable of acting in different positions within a protein (amino terminal, internal, and carboxyl terminal). However, the sequence “IHPYW” forming the basis of DegV1 appears to be relatively uncommon in nature and is not present in other proteins encoded in the *S. cerevisiae* genome. In fact, DegV1 appears to be absent from the currently annotated and available fungal genomes in the Saccharomyces Genome Database^44^. Based on our selection criteria, IHPYW alone is unlikely to be the most potent ER degron. In fact, we were able to demonstrate that just by increasing the length of our short degron, we could improve the degron to be more like that of a full ERAD substrate protein. We expect that in most unfolded proteins, multiple short linear degrons would contribute to effective recognition and degradation by quality control systems.

Even with the limitations, we were able to demonstrate the utility of DegV1 and future degrons by illuminating that degrons can function even in evolutionarily distant species (*S. cerevisiae* and *H. sapiens*). This was somewhat surprising because specific degrons transferred between fungi and animals are not always functionally conserved^45^. It should be noted that the degradation rates of our degron-containing proteins are similar to the rates of other well-characterized ERAD substrates. We interpret these results to mean that DegV1 is degraded in a manner consistent with that of endogenous ERAD substrates. The retrotranslocation process itself appears to function by unfolding, or mostly unfolding, its substrates. Few protein transport systems can transport fully-folded proteins across a lipid bilayer with the notable exceptions being the twin arginine transporter (TAT) system (for review see ^46^) and peroxisomal import machinery^47–51^. However, previous studies in the ERAD field provide conflicting accounts over the ability to transport fully-folded proteins from the ER to cytosol^52–54^. What is certain is that glycosylated ERAD substrate proteins are retrotranslocated across the ER membrane, with N-linked glycans representing a similar steric challenge compared to secondary structure or smaller folded proteins^55^. Because DegV1 allows degradation of fully-folded proteins, it is possible that the proteins are transported in a fully-folded state. Although, it seems unlikely that a single heterodimeric channel^24^ or homodimeric channel^28^ would be able to transport a fully-folded substrate. Perhaps, a substrate could occupy multiple ERAD complexes, could require other unidentified proteins, or could use mechanics similar to the peroxisomal machinery^51^.

However, we favor the idea that the targeted proteins are likely recognized and transported in an unfolded state after ERAD complex engagement. Future studies are needed to identify and characterize the molecular mechanisms that underpin this unexpected observation and it will be important to understand whether the core ERAD machinery, ER chaperones, or perhaps additional unidentified components, are required.

Here, we have created the first tool that can be used to begin to answer these outstanding questions. We’ve identified a genetically encoded sequence that is easily manipulatable for targeting proteins for degradation from a previously unreachable cellular localization. The approach we’ve used will, eventually, enable targeted protein degradation from the ER to target physiologically, or pathophysiologically, relevant proteins. We anticipate that this platform will allow targeting and manipulation of a wide range of previously inaccessible proteins, not only for basic research, but for translational studies and, eventually, biomedical therapies.

## Methods

### Yeast strains and plasmids

Yeast were cultured at 30°C in synthetic complete medium (SC) supplemented with the appropriate amino acids. The *hrd1Δ* and *pdr5Δ* strain used in this study were derivatives of BY4741 (MATa *his3Δ1 leu2Δ0 met15Δ0 ura3Δ0*) or BY4742 (*MATα his3Δ1 leu2Δ0505lys2Δ0 ura3Δ0*). The *hrd1Δpdr5Δ* strain was generated by crossing *hrd1Δ* and *pdr5Δ* strains, sporulating the diploids, and screening the appropriate loci by PCR. For a list of yeast strains used in this study, see Supplemental table S1. For a list of plasmids used in this study, see Supplemental table S2. Plasmids were constructed using restriction enzyme cloning or NEB HiFi assembly. Plasmids used in this study were either centromeric^56^ or custom integrating plasmids^57^. For a list of primers used to generate the pentapeptide library, see Supplemental table S3. Plasmids were transformed into yeast using the LiAc/PEG method^58^. Following transformation into yeast, 3-4 independent transformants were passaged 1-2 times on selection media before using in experiments.

### Mammalian cell culture and transfection

U-2 OS and HEK293 cells were cultured in DMEM containing 4.5 g/liter glucose and L-glutamine, and supplemented with 10% fetal bovine serum (Corning) at 37°C and 5% CO_2_. Cells at 60-80% confluence were transiently transfected with the indicated plasmids using Lipofectamine 2000 (Invitrogen, 11668019) according to manufacturer’s protocols. After 24 hours, cells were split for emetine-chase or chemical treatment assays.

### Mammalian cell lysis

Cells were treated with a translation inhibitor, 50μM emetine (Calbiochem, 324693) for the indicated time periods. For chemical treatments, cells were either mock treated with DMSO, treated with 50 nM bortezomib (APExBIO, A2614), or treated with 1 μM CB5083 (Cayman Chemicals, 19311) for the indicated time periods prior to collection. For combination treatments, cells were pretreated with either bortezomib, or CB5083, for 16 hours prior to emetine chase. The cells were collected and washed once in 1x PBS (10 mM phosphate buffered saline, pH 7.4, 138 mM NaCl, 2.7 mM KCl) before lysis in 50 mM Tris, pH 7.4, 150 mM NaCI, 1% Triton X-100, 1 mM PMSF, and protease inhibitor cocktail for 10-20 minutes at 4°C. The lysates were cleared by centrifugation at 20,000 × g for 10 minutes at 4°C. Protein concentrations were determined using a BCA assay (Thermo Fisher Scientific, 23225). Cell lysates were normalized to the same concentrations in Laemmli sample buffer and heated to 65°C for 5 minutes prior to separation by SDS-PAGE, transferred to a PVDF membrane, immunoblotted with antibodies (anti-HA from Roche, anti-β-actin from Cell Signaling, HRP-linked ECL rabbit-IgG and rat-IgG from Cytiva), and detected by chemiluminescence (ECL Select Western blotting detection reagent, Cytiva) using a ChemiDoc MP (Bio-Rad). For quantification of the immunoblot band intensities, we used ImageLab version 6.1 (Bio-Rad). Band intensities were normalized to β-actin protein in the sample quantified within each lane.

### Yeast pentapeptide library generation

We designed an *in vivo* gap repair strategy for cloning our pentapeptide libraries into the ER-tFT. The plasmid backbone was based on pRS416^59^ and contained a *TDH3* promoter (also known as GPD or GAPDH), the signal sequence from mating factor alpha, mCherry, GFP, an HDEL ER retention signal, and the *CYC1* terminator. To prevent the peptide library from potentially disrupting signal sequence cleavage, two amino acids (Ala and Ser) after the signal sequence cleavage site were left upstream of the library. The final N-terminal amino acid sequence of the ER-tFT library after translocation and signal peptide cleavage is ASXXXXX.

To generate the pentapeptide DNA library fragment with homology arms to ER-tFT, four PCRs were performed (Figure S1C) with Phusion polymerase (New England Biolabs, M0530S). We started by generating a linear DNA template to reduce bias in our PCR reactions. Using pRP01 and primers prRP07 and prRP08, we amplified a 1025 bp fragment with 524 bp upstream (overlapping the *TDH3* promoter and signal sequence) and 495 bp downstream (overlapping mCherry) of the library insertion (EcoRI) site in pRP01. This fragment (fragment 1) was gel-purified (Qiagen, 28104) to remove any residual plasmid and contain only a linear template.

Fragment 2, containing the upstream homology arm, was generated through PCR using fragment 1 (the linear DNA template) with primers prRP10 and prRP29 to amplify a 512 bp fragment, which included homology with both the *TDH3* promoter and signal sequence and contained the random DNA library insertions. Fragment 3, containing the downstream homology arm, was generated in a PCR reaction using fragment 1 (the linear DNA template) from the first PCR reaction with primers prRP09 and prRP51 to generate a 146 bp fragment, which included random DNA in the library position along with homology arms in mCherry. Only 102 bp of homology was included in the mCherry coding region to limit the number of mutations found in mCherry included by homologous recombination. When we included longer homology arms into mCherry, we found that our screening procedure was selective enough to identify mutations in mCherry which could lead to false “unstable” hits (data not shown).

To generate the PCR product for homologous recombination containing overlapping homology arms to the target plasmid (pRP01), we gel-purified fragments 2 and 3 and mixed the DNA in an equimolar ratio. Using primers prRP29 and prRP51, we used 25 PCR cycles to generate a 605 bp fragment 4 (containing the pentapeptide library and homology arms covering 488bp upstream and 111 bp downstream of the library cut site). Fragment 4 was purified and mixed at a 30:1 molar ratio with purified pRP01 (digested with EcoRI) immediately before yeast electroporation.

The library transformation into yeast strain yRB203 was performed as previously described^60^. Cells were grown overnight to stationary phase in YPD media, shaking at 225 rpm and 30°C. An aliquot of the overnight culture was used to inoculate 400 mL of YPD media at 0.3 OD_600_/mL. Cells were grown for approximately 5 hrs until 1.6 OD_600_/mL was reached and collected by centrifugation at 3200 x g for 5 minutes. The cell pellet was washed twice by 200 mL of ice-cold water and once by 200 mL of electroporation buffer (1 M Sorbitol/1 mM CaCl_2_, sterile filtered). The cell pellet was then resuspended in 80 mL of 100 mM LiAc/10 mM DTT, split into two aliquots of 40 mL, and each was incubated in a 250 mL culture flask for 30 minutes at 30°C, shaking at 225 rpm. Next, cells were collected by centrifugation, washed once with 200 mL of ice-cold electroporation buffer, and resuspended to 2.4 mL in electroporation buffer. The cell resuspension was evenly divided into 6 pre-chilled BioRad GenePulser cuvettes (0.2 cm electrode gap) and kept on ice for 10 minutes with DNA. One reaction was used as a no DNA control, one reaction received digested vector only, and four cuvettes received 3 µg of digested vector and 9 µg Fragment 4. Cells were electroporated at 2.5 kV and 25 µF, with time constants varying from 4.0-4.3 milliseconds. Cells were gently transferred from each cuvette into 8 mL of a 1:1 mix of 1M sorbitol:YPD in culture tubes (25m diameter) and incubated at 30°C, with shaking at 220 rpm. After 1 hour, cells were pelleted by centrifugation and inoculated into 1 L SC dropout media (-ura) at 0.2 OD_600_/mL. Dilutions from the electroporated cells were also plated on SC dropout plates and grown for two days at 30°C to determine the library transformation size (approximately 1.2 million for the library described here).

Electroporated cells were grown for ∼18 hours while shaking at 30°C until reaching 0.5 OD_600_/mL. Control strains expressing ER-tFT (pRP01) and KHN-tFT (pRP08) were also cultured in parallel. The strains and library were then treated with DMSO only (Sigma-Aldrich, D2650) or 50 µg/mL cycloheximide (EMD Millipore, 239763) for 2 hours. After treatment, the cells were pelleted, washed once in 1x PBS, resuspended in 1x PBS containing 1 µM Sytox Blue (Invitrogen, S11348), and incubated at 4°C prior to cell sorting.

### Fluorescence-activated cell sorting

Cells were sorted on a MoFlo Astrios Cell Sorter (Beckman Coulter) running Summit software. The instrument was set with a 100 µm tip, 405 nm laser with 448/59 nm bandpass filter, 488 nm laser with 514/20 nm bandpass filter, and 561 nm laser with 620/29 nm bandpass filter. Events were gated to select for yeast cell-sized events, single cells, live cells, and mCherry-GFP positive cells (Figure S1A). The mCherry/GFP ratio for each cell in the final gated population was displayed as a histogram. The ER-tFT and KHN-tFT controls were used to help define where to draw the low mCherry/GFP Ratio (unstable) bin for sorting (Figure S1D). In total, 13 million mCherry-GFP positive events were sorted from the pentapeptide-ER-tFT library, >10 times over the library size. The sorted cells were grown at 30°C with shaking in 5 mL of SC dropout media for 24 hours and expanded to 25 mL cultures overnight. 10 OD_600_ of both the “unstable” bin and unsorted pentapeptide-ER-tFT library were pelleted and flash frozen in liquid nitrogen and stored at -80°C prior to DNA extraction.

### DNA extraction and amplicon sequencing prep

DNA extraction was performed essentially as previously described^34^. The frozen 10 OD_600_ pellets were resuspended in 500 µL of 10 mM K_2_HPO_4_ pH 7.2, 10 mM EDTA, 50 mM 2-mercaptoethanol and incubated with 50 mg/mL zymolyase 100T (AMSBIO) at 37°C for 30-60 minutes until the mixture became clear. 100 µL of lysis buffer (25 mM Tris-HCl pH 7.5, 25 mM EDTA, 2.5% SDS (w/v)) was added and the suspension was incubated at 65°C for 45 minutes. Proteins were precipitated by adding 166 µL of 3M potassium acetate and incubating on ice for 10 minutes. Samples were then centrifuged at 21,000 x g for 10 minutes at 4°C. The supernatant containing DNA was collected and the DNA was precipitated by the addition of 800 µL of 100% ethanol, followed by centrifugation at 21,000 x g for 10 minutes at 4°C. Precipitated DNA was washed with 70% (v/v) ethanol and resuspended in 80 µL of water.

Next, 5 µL of the isolated DNA solution to amplify a 217 bp fragment encompassing the pentapeptide sequences. Partial adapters for Illumina sequencing were added by 25 cycles of PCR using primers prRP37 and prRP38 (annealing temperature of 60°C using Phusion DNA polymerase). PCR products were purified (Qiagen, 28104), normalized to 20 ng/µl using a QuBit 3 (Invitrogen, Q32854), and sent for amplicon sequencing using Genewiz Amplicon-EZ (now Azenta Life Sciences).

### Amplicon-EZ analysis

Sequences from the two Amplicon-EZ samples (unstable bin and input library) were analyzed for quality, trimmed, aligned, and translated by Genewiz (now Azenta Life Sciences). The sequences and translations can be found in Supplementary data, sheets labeled Library Data and Sorted Bin Data. From the translated sequences, the pentapeptide region was isolated and divided into positions #1-5 (P1-P5). The amino acid count was the sum of occurrences for each amino acid at each position (Supplementary data S1, AA Analysis Sheet). We then divided the amino acid frequency at each position for the library or unstable sorted bin by the expected amino acid frequency based on the number of codons that encode a given amino acid. This gave us the relative enrichment of each amino acid at each of the five positions (Figure 1F).

### Flow cytometry based degradation assays

For each experiment, two biological replicates were transferred into SC dropout media in a 96 well plate (Fisherbrand, 12566611) sealed with gas-permeable membranes (Sigma-Aldrich, Z763624) and grown overnight shaking at 1000 rpm at 30°C. Overnight cell density was typically around ∼4-5 OD_600_/mL. In the morning, cells were diluted to 0.2 OD_600_/mL in SC dropout media and grown at 30°C shaking at 1000 rpm for ∼5 hours or until the OD_600_/mL of the cultures was >= 0.5. Cells were pelleted at 3,200 x g and the supernatant was removed by aspiration prior to resuspension in SC dropout media at 2 OD_600_/mL. For experiments using cells treated with 50 µg/mL cycloheximide or 50 µM bortezomib, cells were transferred directly into new 96 well plates (Grenier Bio-One, 650185). To account for slowed growth by these treatments, in the same plates the untreated/DMSO-treated cells were diluted by ⅓ with fresh media. During the treatment periods, cells were incubated at 30°C while shaking at 600 rpm. After treatment, cells were pelleted at 3,200 x g, washed once with 1x PBS, and resuspended in 1x PBS with 1 µM Sytox Blue (Invitrogen). Cells were maintained at 4°C during flow cytometry analysis on a MACSQuant VYB (Miltenyi) running MACSQuantify software (version 2.13.2). Sytox Blue was followed using the 405 nm laser and 452/45 nm emission filters. SuperfastGFP was followed using the 488 nm laser and 452/45 nm emission filters. mCherry was followed using the 561 nm laser 615/20 nm emission filters. Downstream analyses were performed in FlowJo (version 10.7.1) with event gating to select yeast cell-sized events, single cells, live cells, and mCherry-GFP positive cells (Figure S1A). The mCherry/GFP ratio of mCherry-GFP positive cells was calculated in FlowJo and analyzed using a one-way ANOVA and Tukey’s multiple comparisons tests in Graphpad Prism.

### Immunoblotting based degradation assays

Cycloheximide-chase degradation assays were performed as described previously^57,61^ with the following modifications. Starter cultures were grown overnight in SC dropout media while shaking at 30°C. Cultures were diluted to 0.2 OD_600_/mL in SC dropout media and grown for 4 -5 hours to mid-log phase (0.4 -1.0 OD_600_/mL). Cultures were pelleted at 3,200 x g for 5 minutes and resuspended to 2.0 OD_600_/mL in fresh media before treatment with 50 µg/mL cycloheximide. Samples were incubated at 30°C with shaking and, at the indicated time points, shifted to 4°C and collected by centrifugation at 21,000 x g for 5 minutes. The supernatant was removed and cell pellets were incubated on dry ice prior to storage at -80°C or cell lysis. For experiments with bortezomib treatment, cells were pre-treated with 50 µM bortezomib for 15 minutes prior to cycloheximide addition.

Cells were resuspended in SUME lysis buffer (1% SDS, 8M urea, 10 mM MOPS, pH 6.8, 10mM EDTA)^62^ at 20 OD_600_/mL with acid-washed glass beads (0.1 mm, Bio-Spec). Cells were vortexed for 2 minutes and an equal volume of sample buffer (4% SDS, 8M urea, 125 mM Tris pH 6.8, 10% β-mercaptoethanol, 0.02 % bromophenol blue) was added and briefly vortexed. The samples were incubated at 65°C for 5 minutes, separated by SDS-PAGE, transferred to a PVDF membrane, immunoblotted with antibodies (anti-GFP from Genscript, anti-DYKDDDK from Genscript, anti-HA from Roche, anti-V5 from Genscript, HRP-linked ECL rabbit-IgG and mouse-IgG from Cytiva, Goat anti-Mouse IgG Alexa800 from Invitrogen), and detected by chemiluminescence (ECL Select Western blotting detection reagent, Cytiva) or by Dylight800 fluorescence using a ChemiDoc MP (Bio-Rad). For quantification of the immunoblot band intensities, we used ImageLab version 6.1 (Bio-Rad). Band intensities were normalized to total protein in the sample quantified within each lane using Stain-Free Dye Imaging (Bio-Rad).

## Acknowledgements

We would like to thank Ling Qi for sharing the Hrd1-deficient HEK293 cells. We would also like to thank Andrew Folkmann and Kaushik Ragunathan for their critical reading of the manuscript, Basila Moochickal Assainar for help with the fluorescence microscopy, and other members of the Baldridge lab for their thoughtful discussion and comments regarding this work. R.S. (previously R. Plumb) was supported by an NIH/NIGMS Award (5F32GM136020) and D.D. was supported by the NIH Cellular and Molecular Biology Training Grant (T32-GM007315). In addition, this work was supported by an NIH/NIGMS Award (R35GM128592 to R.D.B.).

## Author contributions

R.S. conceptualized the project, developed the methodology, performed the investigation, analyzed the data, visualized the data, wrote the original draft, and reviewed/edited the final manuscript. J.H. developed the methodology, performed the investigation, analyzed the data, visualized the data, and reviewed/edited the final manuscript. D.D. performed the investigation, analyzed the data, visualized the data, and reviewed/edited the final manuscript. R.D.B. conceptualized the project, performed the investigation, analyzed the data, visualized the data, wrote the original draft, reviewed/edited the final manuscript, and supervised the study.

## Materials Availability Statement

Further information and requests for resources should be directed to and will be fulfilled by the lead contact, Ryan Baldridge.

## Declaration of Interests

R.D.B. and R.S. have filed a provisional patent application covering the use of this technology.

## Supplemental Information

**Figure S1.**
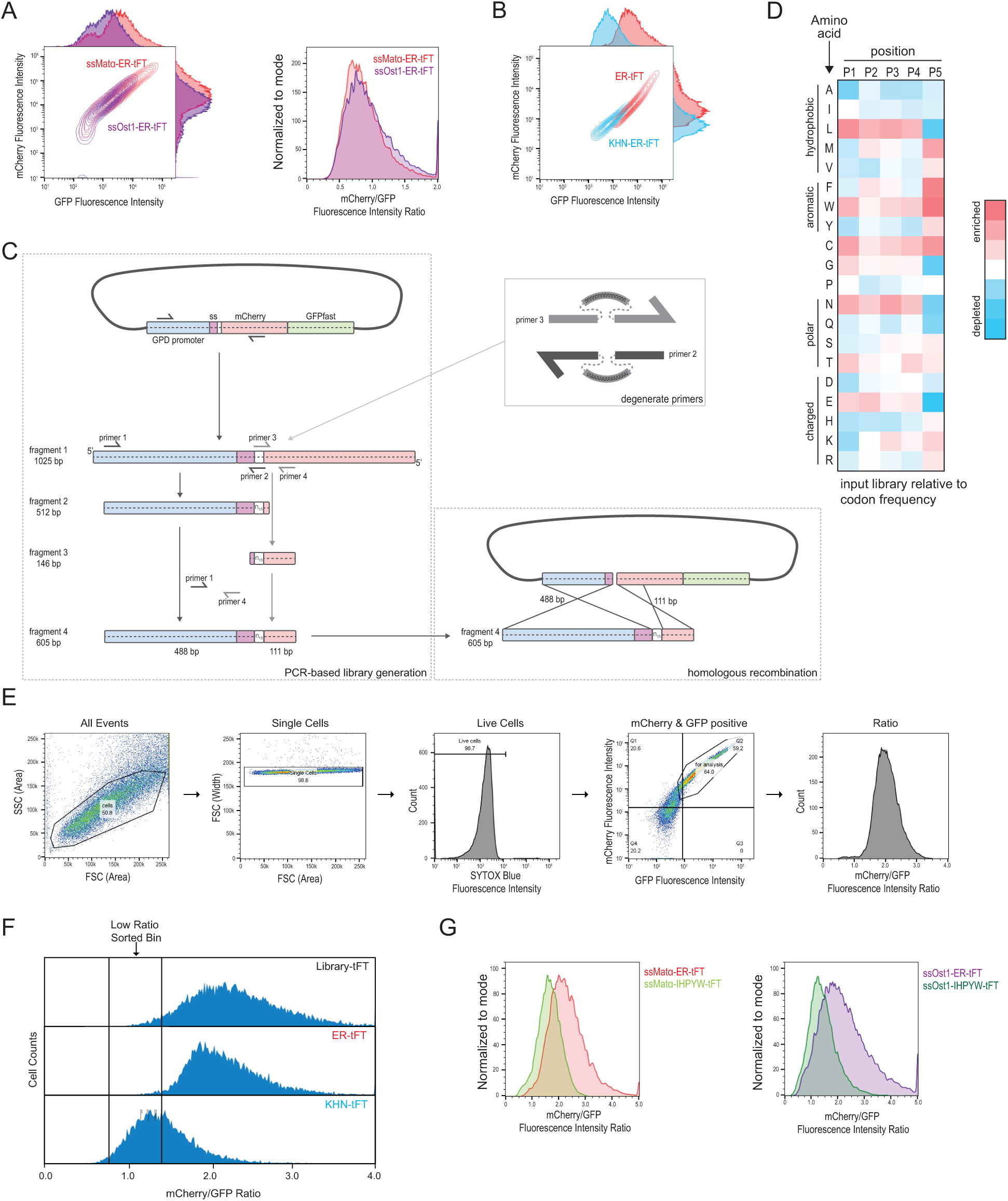
(A) The tandem fluorescent timer (tFT) was targeted to the ER with two different signal peptides, either that of mating factor alpha (Matα) or of Ost1. The GFP and mCherry fluorescence were displayed as a contour plot (left panel). The mCherry/GFP fluorescence intensity ratio of each cell was calculated displayed as a histogram (right panel). Based on the superior brightness of cells expressing the mating factor alpha signal sequence, we selected this signal peptide for further experimentation. (B) Flow cytometry of yeast strains expressing an ER-tFT and ERAD-substrate KHN-tFT treated with cycloheximide for 2 hours. (C) Schematic of the PCR-mediated library generation (left) using degenerate primers (upper right) and homologous recombination in yeast (lower right). (D) Heatmap of input library amino acid enrichments at each position displayed relative to codon frequency. (E) Gating strategy for analyzing single cells by flow cytometry. (F) Sorting bins defined relative to ER-tFT and KHN-tFT. (G) Comparison of mCherry/GFP ratios for alternate signal sequences on ER-tFT and IHPYW-tFT.

**Figure S2.**
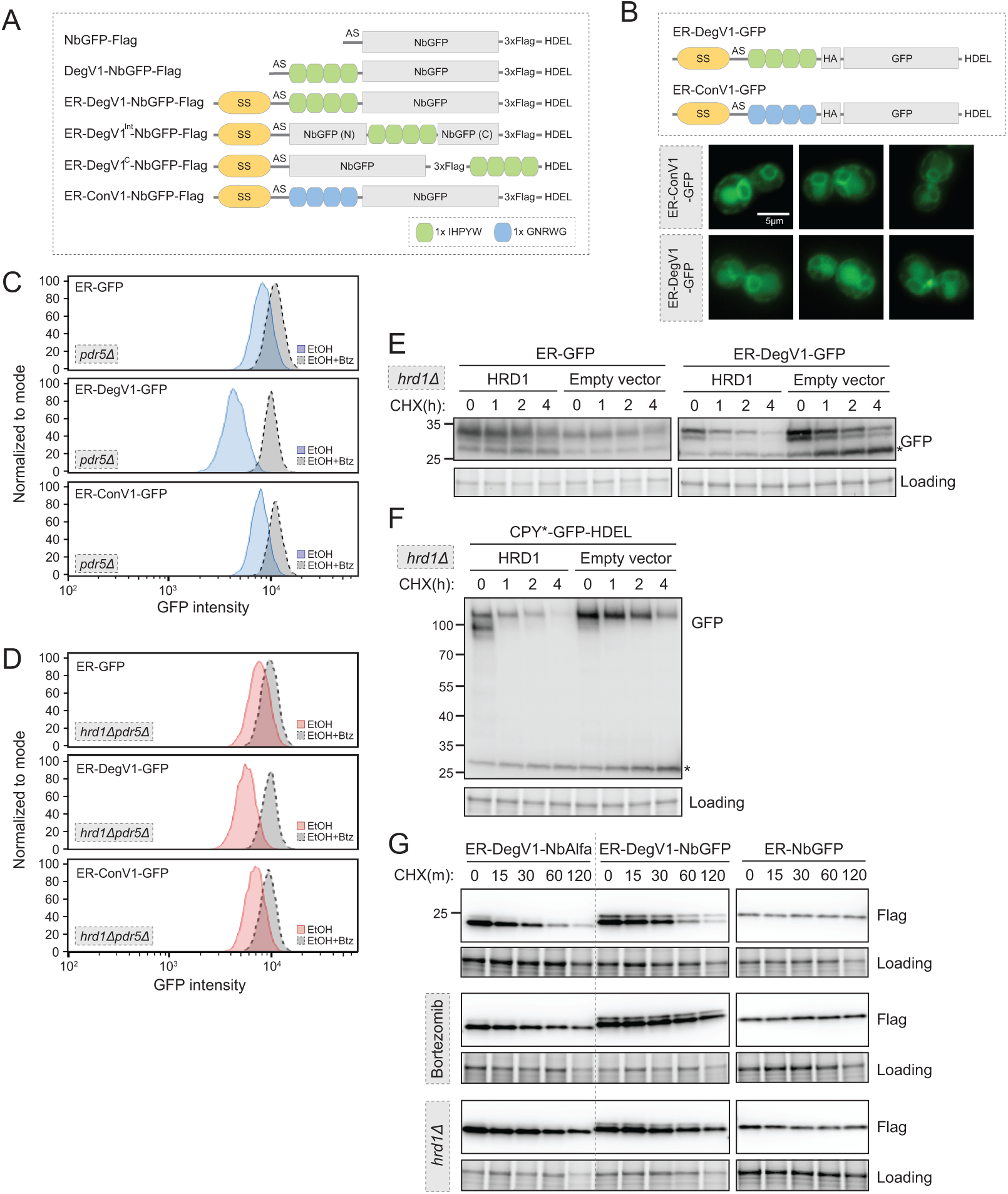
(A) Linear diagram of degradation constructs used in this work. SS, mating factor alpha signal sequence; AS, alanine and serine dipeptide linker; NbGFP, LaG16 anti-GFP nanobody; HDEL, ER retention signal. (B) Localization of ER-targeted GFP proteins (ER-ConV1-GFP and ER-DegV1-GFP) in wild-type yeast cells. Proteins were observed in both the ER and vacuolar localization. The scale bar is 5 µm. (C) The degradation of ER-targeted GFP proteins (ER-GFP (top), ER-DegV1-GFP (middle), or ER-ConV1-GFP (bottom)) were analyzed by flow cytometry following either ethanol (EtOH) or with bortezomib (Btz) for 2 hours. (D) As in (C), but in a *hrd1Δpdr5Δ* strain. (E) Degradation of ER-targeted GFP with an ER retention signal (HDEL) with, or without DegV1 was followed in a *hrd1Δ* strain complemented with either an empty vector or with Hrd1 on a centromeric plasmid. Using a cycloheximide chase and immunoblotting, we found in the absence of Hrd1, we observe transport of the proteins to the vacuole, where free GFP accumulates. The asterisk indicates the vacuolar localized GFP fragment. (F) As in (E), except with CPY*-GFP-HDEL. The asterisk indicates the vacuolar localized GFP fragment. (G) Degradation of ER-DegV1-NbAlfa, ER-DegV1-NbGFP or ER-NbGFP was followed using a cycloheximide chase in the presence or absence of bortezomib (Btz) in a *pdr5Δ* or *hrd1Δpdr5Δ* strain. All panels in this figure are representative of at least three independent biological replicates.

**Figure S3.**
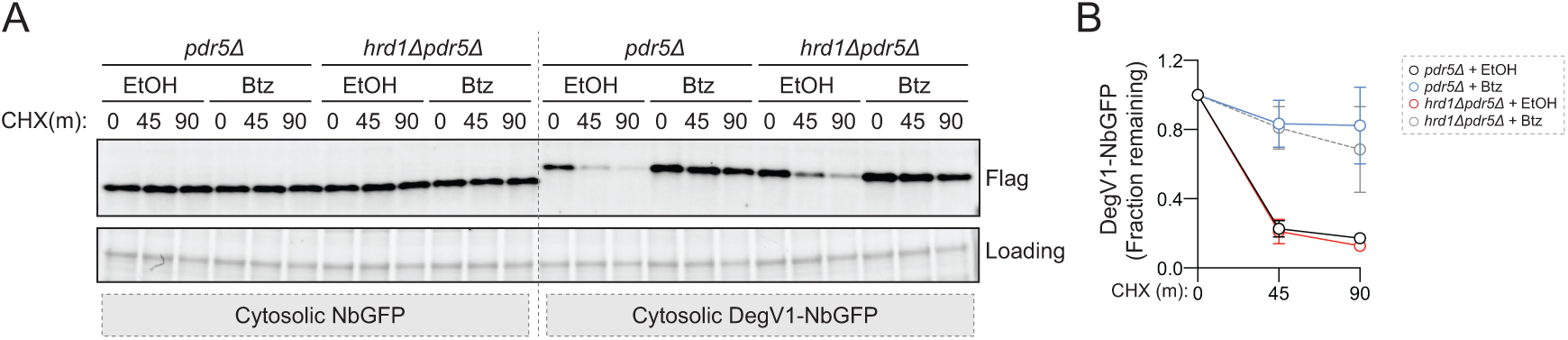
(A) Degradation of a cytosolically-localized anti-GFP nanobody (cytosolic NbGFP) or with DegV1 (cytosolic DegV1-NbGFP) was followed in *pdr5Δ or hrd1Δpdr5Δ* strains with, or without bortezomib (Btz) using a cycloheximide (CHX) chase. Note that Hrd1 is not required for the degradation of cytosolic DegV1-NbGFP. (C) Quantification of three independent biological replicates from (A). Error bars represent the standard deviation.

## Notes

### Summary of Updates

There are minor revisions to the text.

## References

1. Collins, G.A., and Goldberg, A.L. (2017). The Logic of the 26S Proteasome. Cell 169, 792–806. 10.1016/j.cell.2017.04.023.

2. Clague, M.J., Heride, C., and Urbé, S. (2015). The demographics of the ubiquitin system. Trends Cell Biol. 25, 417–426. 10.1016/j.tcb.2015.03.002.

3. Bachmair, A., Finley, D., and Varshavsky, A. (1986). In Vivo Half-Life of a Protein Is a Function of Its Amino-Terminal Residue. Science 234, 179–186. 10.1126/science.3018930.

4. Varshavsky, A. (2011). The N-end rule pathway and regulation by proteolysis. Protein Science 20, 1298–1345. 10.1002/pro.666.

5. Varshavsky, A. (2019). N-degron and C-degron pathways of protein degradation. Proc. Natl. Acad. Sci. U.S.A. 116, 358–366. 10.1073/pnas.1816596116.

6. Geffen, Y., Appleboim, A., Gardner, Richard G., Friedman, N., Sadeh, R., and Ravid, T. (2016). Mapping the Landscape of a Eukaryotic Degronome. Mol. Cell 63, 1055–1065. 10.1016/j.molcel.2016.08.005.

7. Mashahreh, B., Armony, S., Johansson, K.E., Chappleboim, A., Friedman, N., Gardner, R.G., Hartmann-Petersen, R., Lindorff-Larsen, K., and Ravid, T. (2022). Conserved degronome features governing quality control associated proteolysis. Nat Commun 13, 7588. 10.1038/s41467-022-35298-y.

8. Koren, I., Timms, R.T., Kula, T., Xu, Q., Li, M.Z., and Elledge, S.J. (2018). The Eukaryotic Proteome Is Shaped by E3 Ubiquitin Ligases Targeting C-Terminal Degrons. Cell 173, 1622–1635.e1614. 10.1016/j.cell.2018.04.028.

9. Lin, H.-C., Yeh, C.-W., Chen, Y.-F., Lee, T.-T., Hsieh, P.-Y., Rusnac, D.V., Lin, S.-Y., Elledge, S.J., Zheng, N., and Yen, H.-C.S. (2018). C-Terminal End-Directed Protein Elimination by CRL2 Ubiquitin Ligases. Mol. Cell 70, 602–613.e603. 10.1016/j.molcel.2018.04.006.

10. Timms, R.T., and Koren, I. (2020). Tying up loose ends: the N-degron and C-degron pathways of protein degradation. Biochem. Soc. Trans. 48, 1557–1567. 10.1042/bst20191094.

11. Samarasinghe, K.T.G., and Crews, C.M. (2021). Targeted protein degradation: A promise for undruggable proteins. Cell Chemical Biology 28, 934–951. 10.1016/j.chembiol.2021.04.011.

12. Christianson, J.C., and Carvalho, P. (2022). Order through destruction: how ER-associated protein degradation contributes to organelle homeostasis. EMBO J. 41, e109845. 10.15252/embj.2021109845.

13. Carvalho, P., Goder, V., and Rapoport, T.A. (2006). Distinct ubiquitin-ligase complexes define convergent pathways for the degradation of ER proteins. Cell 126, 361–373. 10.1016/j.cell.2006.05.043.

14. Denic, V., Quan, E.M., and Weissman, J.S. (2006). A luminal surveillance complex that selects misfolded glycoproteins for ER-associated degradation. Cell 126, 349–359. 10.1016/j.cell.2006.05.045.

15. Foresti, O., Rodriguez-Vaello, V., Funaya, C., and Carvalho, P. (2014). Quality control of inner nuclear membrane proteins by the Asi complex. Science 346, 751–755. 10.1126/science.1255638.

16. Khmelinskii, A., Blaszczak, E., Pantazopoulou, M., Fischer, B., Omnus, D.J., Le Dez, G., Brossard, A., Gunnarsson, A., Barry, J.D., Meurer, M., et al. (2014). Protein quality control at the inner nuclear membrane. Nature 516, 410–413. 10.1038/nature14096.

17. Huyer, G., Piluek, W.F., Fansler, Z., Kreft, S.G., Hochstrasser, M., Brodsky, J.L., and Michaelis, S. (2004). Distinct machinery is required in Saccharomyces cerevisiae for the endoplasmic reticulum-associated degradation of a multispanning membrane protein and a soluble luminal protein. J. Biol. Chem. 279, 38369–38378. 10.1074/jbc.M402468200.

18. Vashist, S., and Ng, D.T.W. (2004). Misfolded proteins are sorted by a sequential checkpoint mechanism of ER quality control. J. Cell Biol. 165, 41–52. 10.1083/jcb.200309132.

19. Bays, N.W., Wilhovsky, S.K., Goradia, A., Hodgkiss-Harlow, K., and Hampton, R.Y. (2001). HRD4/NPL4 is required for the proteasomal processing of ubiquitinated ER proteins. Mol. Biol. Cell 12, 4114–4128. 10.1091/mbc.12.12.4114.

20. Jarosch, E., Taxis, C., Volkwein, C., Bordallo, J., Finley, D., Wolf, D.H., and Sommer, T. (2002). Protein dislocation from the ER requires polyubiquitination and the AAA-ATPase Cdc48. Nat. Cell Biol. 4, 134–139. 10.1038/ncb746.

21. Rabinovich, E., Kerem, A., Fröhlich, K.-U., Diamant, N., and Bar-Nun, S. (2002). AAA-ATPase p97/Cdc48p, a cytosolic chaperone required for endoplasmic reticulum-associated protein degradation. Mol. Cell. Biol. 22, 626–634. 10.1128/MCB.22.2.626-634.2002.

22. Ye, Y., Meyer, H.H., and Rapoport, T.A. (2001). The AAA ATPase Cdc48/p97 and its partners transport proteins from the ER into the cytosol. Nature 414, 652–656. 10.1038/414652a.

23. Mehnert, M., Sommer, T., and Jarosch, E. (2014). Der1 promotes movement of misfolded proteins through the endoplasmic reticulum membrane. Nat. Cell Biol. 16, 77–86. 10.1038/ncb2882.

24. Wu, X., Siggel, M., Ovchinnikov, S., Mi, W., Svetlov, V., Nudler, E., Liao, M., Hummer, G., and Rapoport, T.A. (2020). Structural basis of ER-associated protein degradation mediated by the Hrd1 ubiquitin ligase complex. Science 368, eaaz2449. 10.1126/science.aaz2449.

25. Pisa, R., and Rapoport, T.A. (2022). Disulfide-crosslink analysis of the ubiquitin ligase Hrd1 complex during endoplasmic reticulum-associated protein degradation. J. Biol. Chem., 102373. 10.1016/j.jbc.2022.102373.

26. Baldridge, R.D., and Rapoport, T.A. (2016). Autoubiquitination of the Hrd1 Ligase Triggers Protein Retrotranslocation in ERAD. Cell 166, 394–407. 10.1016/j.cell.2016.05.048.

27. Carvalho, P., Stanley, A.M., and Rapoport, T.A. (2010). Retrotranslocation of a misfolded luminal ER protein by the ubiquitin-ligase Hrd1p. Cell 143, 579–591. 10.1016/j.cell.2010.10.028.

28. Schoebel, S., Mi, W., Stein, A., Ovchinnikov, S., Pavlovicz, R., DiMaio, F., Baker, D., Chambers, M.G., Su, H., Li, D., et al. (2017). Cryo-EM structure of the protein-conducting ERAD channel Hrd1 in complex with Hrd3. Nature 548, 352–355. 10.1038/nature23314.

29. Stein, A., Ruggiano, A., Carvalho, P., and Rapoport, T.A. (2014). Key steps in ERAD of luminal ER proteins reconstituted with purified components. Cell 158, 1375–1388. 10.1016/j.cell.2014.07.050.

30. Needham, P.G., Guerriero, C.J., and Brodsky, J.L. (2019). Chaperoning Endoplasmic Reticulum–Associated Degradation (ERAD) and Protein Conformational Diseases. Cold Spring Harb Perspect Biol 11. 10.1101/cshperspect.a033928.

31. Fisher, A.C., and DeLisa, M.P. (2008). Laboratory Evolution of Fast-Folding Green Fluorescent Protein Using Secretory Pathway Quality Control. PLoS ONE 3, e2351. 10.1371/journal.pone.0002351.

32. Shaner, N.C., Campbell, R.E., Steinbach, P.A., Giepmans, B.N.G., Palmer, A.E., and Tsien, R.Y. (2004). Improved monomeric red, orange and yellow fluorescent proteins derived from Discosoma sp. red fluorescent protein. Nat. Biotechnol. 22, 1567–1572. 10.1038/nbt1037.

33. Khmelinskii, A., Keller, P.J., Bartosik, A., Meurer, M., Barry, J.D., Mardin, B.R., Kaufmann, A., Trautmann, S., Wachsmuth, M., Pereira, G., et al. (2012). Tandem fluorescent protein timers for in vivo analysis of protein dynamics. Nat. Biotechnol. 30, 708–714. 10.1038/nbt.2281.

34. Kats, I., Khmelinskii, A., Kschonsak, M., Huber, F., Knieß, R.A., Bartosik, A., and Knop, M. (2018). Mapping Degradation Signals and Pathways in a Eukaryotic N-terminome. Mol. Cell 70, 488–501.e485. 10.1016/j.molcel.2018.03.033.

35. Vashist, S., Kim, W., Belden, W.J., Spear, E.D., Barlowe, C., and Ng, D.T. (2001). Distinct retrieval and retention mechanisms are required for the quality control of endoplasmic reticulum protein folding. J. Cell Biol. 155, 355–368. 10.1083/jcb.200106123.

36. Fridy, P.C., Li, Y., Keegan, S., Thompson, M.K., Nudelman, I., Scheid, J.F., Oeffinger, M., Nussenzweig, M.C., Fenyö, D., Chait, B.T., and Rout, M.P. (2014). A robust pipeline for rapid production of versatile nanobody repertoires. Nat. Methods 11, 1253–1260. 10.1038/nmeth.3170.

37. Bordallo, J., Plemper, R.K., Finger, A., and Wolf, D.H. (1998). Der3p/Hrd1p is required for endoplasmic reticulum-associated degradation of misfolded lumenal and integral membrane proteins. Mol. Biol. Cell 9, 209–222. 10.1091/mbc.9.1.209.

38. Azuma, M., Levinson, J.N., Pagé, N., and Bussey, H. (2002). Saccharomyces cerevisiae Big1p, a putative endoplasmic reticulum membrane protein required for normal levels of cell wall β-1,6-glucan. Yeast 19, 783–793. 10.1002/yea.873.

39. Toke, D.A., and Martin, C.E. (1996). Isolation and characterization of a gene affecting fatty acid elongation in Saccharomyces cerevisiae. J. Biol. Chem. 271, 18413–18422. 10.1074/jbc.271.31.18413.

40. Nie, L., Pascoa, T.C., Pike, A.C.W., Bushell, S.R., Quigley, A., Ruda, G.F., Chu, A., Cole, V., Speedman, D., Moreira, T., et al. (2021). The structural basis of fatty acid elongation by the ELOVL elongases. Nature Structural & Molecular Biology 28, 512–520. 10.1038/s41594-021-00605-6.

41. Shaner, N.C., Lambert, G.G., Chammas, A., Ni, Y., Cranfill, P.J., Baird, M.A., Sell, B.R., Allen, J.R., Day, R.N., Israelsson, M., et al. (2013). A bright monomeric green fluorescent protein derived from Branchiostoma lanceolatum. Nat. Methods 10, 407–409. 10.1038/nmeth.2413.

42. Anderson, D.J., Le Moigne, R., Djakovic, S., Kumar, B., Rice, J., Wong, S., Wang, J., Yao, B., Valle, E., von Soly, S.K., et al. (2015). Targeting the AAA ATPase p97 as an approach to treat cancer through disruption of protein homeostasis. Cancer cell 28, 653–665. 10.1016/j.ccell.2015.10.002.

43. Shi, G., Somlo, D., Kim, G.H., Prescianotto-Baschong, C., Sun, S., Beuret, N., Long, Q., Rutishauser, J., Arvan, P., Spiess, M., and Qi, L. (2017). ER-associated degradation is required for vasopressin prohormone processing and systemic water homeostasis. J. Clin. Invest. 127. 10.1172/JCI94771.

44. Cherry, J.M., Hong, E.L., Amundsen, C., Balakrishnan, R., Binkley, G., Chan, E.T., Christie, K.R., Costanzo, M.C., Dwight, S.S., Engel, S.R., et al. (2012). Saccharomyces Genome Database: the genomics resource of budding yeast. Nucleic Acids Res 40, D700–705. 10.1093/nar/gkr1029.

45. Timms, R.T., Zhang, Z., Rhee, D.Y., Harper, J.W., Koren, I., and Elledge, S.J. (2019). A glycine-specific N-degron pathway mediates the quality control of protein N-myristoylation. Science 365, eaaw4912. 10.1126/science.aaw4912.

46. Berks, B.C. (2015). The Twin-Arginine Protein Translocation Pathway. Annu. Rev. Biochem. 84, 843–864. 10.1146/annurev-biochem-060614-034251.

47. McNew, J.A., and Goodman, J.M. (1994). An oligomeric protein is imported into peroxisomes in vivo. Journal of Cell Biology 127, 1245–1257. 10.1083/jcb.127.5.1245.

48. Glover, J.R., Andrews, D.W., and Rachubinski, R.A. (1994). Saccharomyces cerevisiae peroxisomal thiolase is imported as a dimer. Proc. Natl. Acad. Sci. U.S.A. 91, 10541–10545. 10.1073/pnas.91.22.10541.

49. Romano, F.B., Blok, N.B., and Rapoport, T.A. (2019). Peroxisome protein import recapitulated in Xenopus egg extracts. J. Cell Biol. 218, 2021–2034. 10.1083/jcb.201901152.

50. Skowyra, M.L., and Rapoport, T.A. (2022). PEX5 translocation into and out of peroxisomes drives matrix protein import. Mol. Cell 82, 3209–3225.e3207. 10.1016/j.molcel.2022.07.004.

51. Gao, Y., Skowyra, M.L., Feng, P., and Rapoport, T.A. (2022). Protein import into peroxisomes occurs through a nuclear pore-like phase. Science 378, eadf3971. 10.1126/science.adf3971.

52. Tirosh, B., Furman, M.H., Tortorella, D., and Ploegh, H.L. (2003). Protein Unfolding Is Not a Prerequisite for Endoplasmic Reticulum-to-Cytosol Dislocation. J. Biol. Chem. 278, 6664–6672. 10.1074/jbc.M210158200.

53. Bhamidipati, A., Denic, V., Quan, E.M., and Weissman, J.S. (2005). Exploration of the topological requirements of ERAD identifies Yos9p as a lectin sensor of misfolded glycoproteins in the ER lumen. Mol. Cell 19, 741–751. 10.1016/j.molcel.2005.07.027.

54. Shi, J., Hu, X., Guo, Y., Wang, L., Ji, J., Li, J., and Zhang, Z.-R. (2019). A technique for delineating the unfolding requirements for substrate entry into retrotranslocons during endoplasmic reticulum-associated degradation. J. Biol. Chem. 294, 20084–20096. 10.1074/jbc.RA119.010019.

55. Grotzke, J.E., Lu, Q., and Cresswell, P. (2013). Deglycosylation-dependent fluorescent proteins provide unique tools for the study of ER-associated degradation. Proc. Natl. Acad. Sci. U.S.A. 110, 3393–3398. 10.1073/pnas.1300328110.

56. Sikorski, R.S., and Hieter, P. (1989). A System of Shuttle Vectors and Yeast Host Strains Designed for Efficient Manipulation of DNA in Saccharomyces ceravisiae. Genetics 122, 19–27.

57. Hwang, J., Peterson, B.G., Knupp, J., and Baldridge, R.D. (2023). The ERAD system is restricted by elevated ceramides. Science Advances 9, eadd8579. 10.1126/sciadv.add8579.

58. Gietz, R.D., and Schiestl, R.H. (2007). High-efficiency yeast transformation using the LiAc/SS carrier DNA/PEG method. Nat Protoc 2, 31–34. 10.1038/nprot.2007.13.

59. Christianson, T.W., Sikorski, R.S., Dante, M., Shero, J.H., and Hieter, P. (1992). Multifunctional yeast high-copy-number shuttle vectors. Gene 110, 119. 10.1016/0378-1119(92)90454-w.

60. Benatuil, L., Perez, J.M., Belk, J., and Hsieh, C.-M. (2010). An improved yeast transformation method for the generation of very large human antibody libraries. Protein Engineering, Design and Selection 23, 155–159. 10.1093/protein/gzq002.

61. Peterson, B.G., Hwang, J., Russ, J.E., Schroeder, J.W., Freddolino, P.L., and Baldridge, R.D. (2023). Deep mutational scanning highlights a role for cytosolic regions in Hrd1 function. Cell Rep. 42. 10.1016/j.celrep.2023.113451.

62. Gardner, R., Cronin, S., Leader, B., Rine, J., and Hampton, R. (1998). Sequence determinants for regulated degradation of yeast 3-hydroxy-3-methylglutaryl-CoA reductase, an integral endoplasmic reticulum membrane protein. Mol. Biol. Cell 9, 2611–2626. 10.1091/mbc.9.9.2611.

